# Kinetics of radiation-induced DNA double-strand breaks through coarse-grained simulations

**DOI:** 10.1101/2022.07.03.498607

**Authors:** Manuel Micheloni, Lorenzo Petrolli, Gianluca Lattanzi, Raffaello Potestio

## Abstract

Double-strand breaks (DSBs), i.e. the covalent cut of the DNA backbone over both strands, are a detrimental outcome of cell irradiation, bearing chromosomal aberrations and leading to cell apoptosis. In the early stages of the evolution of a DSB, the disruption of the residual interactions between the DNA moieties drives the fracture of the helical layout; in spite of its biological significance, the details of this process are still largely uncertain. Here, we address the mechanical rupture of DNA by DSBs *via* coarse-grained molecular dynamics simulations: the setup involves a 3855-bp DNA filament and diverse DSB motifs, i.e. within a range of distances between strand breaks (or DSB distance). By employing a coarse-grained model of DNA, we access the molecular details and characteristic timescales of the rupturing process. A sequence-nonspecific, linear correlation is observed between the DSB distance and the internal energy contribution to the disruption of the residual (Watson-Crick and stacking) contacts between DNA moieties, which is seemingly driven by an abrupt, cooperative process. Moreover, we infer an exponential dependence of the characteristic rupture times on the DSB distances, which we associate to an Arrhenius law of thermally-activated processes. This work lays the foundations of a detailed, mechanistic assessment of DSBs *in silico*, as a benchmark to both numerical simulations and data from single molecule experiments.

## 1. INTRODUCTION

DNA holds the hereditary information required by cells to replicate and carry out their biological activity. Cells, however, are constantly threatened by a variety of toxic agents, from both *endogenous processes*, i.e. naturally associated with a cell life-cycle (e.g. as by-products of metabolic stresses [1] and DNA replication [2]) and *exogenous vectors*, such as chemicals and ionizing radiations (IR) [3, 4]. In fact, IR accounts for a plethora of chemical and mechanical lesions of the DNA molecule, involving a wide range of space and time scales, which are mediated by a cascade of inelastic collisions and radiolytic reactions [5]. Amongst the diverse radiation-enforced DNA lesions, however, the disruption of the phosphoester backbone over both DNA strands, or double-strand break (DSB), is arguably the most toxic [3]. DSBs account for major cytotoxic effects, and critically so at sites of highly-condensed lesions within localized DNA volumes [6, 7]. Indeed, lesion clustering is a striking lethal feature of IR at high linear energy transfer regimes, i.e. associated with a massive release of energy at the microscale [8, 9].

From a radiation biology perspective, a key observable *via* the irradiation of cell-lines *in vitro* is defined by the cell survival ratio *S* upon the radiation dose *D* -the latter being the energy deposited per unit of mass. A simplified, yet effective, theoretical framework that properly describes *S* as a function of *D* is the so-called *linear-quadratic* model whereby 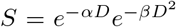: here, *α* and *β* are associated with the cell radiosensitivity and depend on a variety of factors, namely the cell type, the stage of the cell cycle, the dose fractionation regime, the radition quality and energy, and so on [10]. While *α* and *β* are empirically defined, several phenomenological [11] and microdosimetric models [12, 13] have attempted at establishing a mechanistic foundation of the cell survival, relying upon a microscopic description of the layout of the events of energy deposition about a radiation carrier. In this concern, Monte-Carlo track structure (MCTS) codes have been widely employed to characterize *in silico* the evolution of a radiation field as a stochastic sequence of (elastic, inelastic) collisions with a medium -such as a water continuum, in a cell-like scenario -at a resolution level that is hardly accessible by conventional experimental techniques [14–16]. Effective criteria have thus been established to associate the local distribution of physical and chemical processes (or the radiation *track*) with the early occurrence of ‘lethal’ DNA lesions, such as double-strand breaks, within significant sub-cellular volumes [17–19].

Overall, these techniques depict the irradiation of cells *via* a statistically robust, mean field approach, as multiple replicas of the track are averaged. However, the theoretical framework underneath MCTS codes overlooks the dynamical detail to characterize the subsequent evolution of DSBs at the molecular level. In fact, according to a widely-accepted paradigm, double-strand break motifs are implicitly deemed to be effective, achieving a non-reversible rupture of the DNA backbone, within an arbitrary distance of the two strand nicks [17, 18, 20]. Yet, it was earlier inferred that the distribution of DSB motifs is likely associated with a radiation quality, i.e. double-strand breaks occur with a different likelihood upon the distance between cuts [21]. Moreover, the disruption of the residual contact interface holding the DNA moieties should arguably contribute a unique (thermodynamic, kinetic) bias on the local thermal and mechanical stability of DNA varying upon the DSB motif.

A complementary insight might thus be offered by molecular dynamics (MD) techniques, evolving the transient dynamics of DSBs over single DNA molecules. Recently, Landuzzi and co-workers [22] have assessed the thermal and forceful fracture of DNA by DSBs on a 31-bp DNA sequence depicting a short DNA *linker* (i.e. the exposed, linear DNA stretch between nucleosomes in a chromatin fiber) by all-atom MD. Notably, they inferred that the direct observation of the rupture of DNA by DSB motifs beyond a distance of 3 bp is unlikely within the time scales associated with the activity of the DNA damage repair machinery -yet, it is broadly acknowledged that atomistic force fields of nucleic acids might over-estimate the strength of the non-covalent forces holding the DNA helix (such as *π*-stacking contacts) [23, 24]. Moreover, atomistic MD is bound by a significant numerical overhead, so that the accessible dynamics of biological systems is restricted to a few microseconds and/or a few replicas. These limitations may be circumvented by exploiting coarse-grained (CG) models [25–27], whereby groups of atoms are specifically mapped into a lower-resolution description of the system: the CG sites interact by means of effective potentials, defined by either *top-down* or *bottom-up* approaches [24, 28].

In this work, we employ the oxDNA CG [27] force field to assess the rupturing process of DNA driven by double-strand break lesions. We enforce the twofold cuts of the DNA backbone involving no local base modifications nor deletions (or T1-DSBs as described by Schipler and Iliakis [3]) on a 3855-bp DNA molecule, kept at varied end-to-end distances by fixed forces. Here, DSBs are described within a simplified, effective framework, by removing the covalent interactions between adjacent CG nucleotides, varying the distance between strand breaks (0 to 3 bp). The setup resembles an optical tweezers scenario, the DNA being adsorbed and/or fixed at one end of the molecule and slightly stretched by a constant force regime at the other end, taking advantage of an established (experimental, theoretical) framework in single molecule experiments on nucleic acids [29, 30]. We thus perform extensive MD simulations, tracking the transient dynamics of the fractured DNA moieties and the kinetics of the subsequent rupturing process, which is driven by the disruption of all residual contacts holding the helical framework. The trend we observe tallies with a depiction of the DNA rupturing by DSBs as a cooperative, thermally-activated process, characterized by an exponential increase of the average rupture times with the distance between strand breaks. We thus discuss our observations in terms of an Arrhenius law, and propose a protocol to rescale the times of the CG dynamics by the (experimental, *in silico*) diffusion coefficients of freely-diffusing DNA.

## II. MATERIALS AND METHODS

### A. System model and force field

We simulated a 3855-bp, double-stranded DNA sequence in the B conformation (see Section **1** of the Supplementary Material), adapted from the optical tweezing experimental setup of Wang and co-workers [31]. We employed the oxDNA2 CG force field, which has been acknowledged to faithfully reproduce both the thermodynamic and the mechanical features of the DNA double helix [32]. The earliest releases of the model [27, 33] take into account temperature-dependent, sequence-specific stacking interactions, and Watson-Crick hydrogen bonding between nucleotides. oxDNA2 improves on the original model, effectively involving features of the major and minor grooves of B-DNA, and salt-dependent modulations of the force field *via* Debye-Hckel interactions, thereby refining the behaviour of DNA in implicit solvent and diverse concentrations of monovalent salt, down to a physiological environment [32]. Moreover, the level of resolution associated with oxDNA2 is adequate to detail the dynamical features of DSB lesions while lowering the numerical overhead of the atomistic force fields, hence covering timeframes that are biologically significant [24, 34]. It is worth noting that, while the oxDNA2 force field is benchmarked upon experimental, equilibrium properties of DNA, we hereby characterize transient, dynamical processes - so that its validity has to be verified *a posteriori*.

### B. Molecular dynamics simulations

MD simulations were carried out operating on the LAMMPS software platform [35]. The dynamics of the DNA molecules was assessed in the NVT ensemble at *T* = 310 K and a monovalent salt concentration of 0.15 M, matching a physiological environment. All simulations were performed employing the rigid-body Langevin-type integrator (Langevin C) [36], which solves the equations of motion accounting for an implicit solvent background. Indeed, this integrator has shown enhanced performance, in contrast to the standard LAMMPS Langevin algorithm for DNA simulations, while allowing a stable integration of the equations of motion at larger time steps [34, 37] - which we set to Δ*t* = 5 *×* 10^−3^ *τ*, together with a damping coefficient of the Langevin thermostat *ζ* = 2.5 *τ*.

Thus, we carried out a step-wise thermalization protocol (see Section **2** of the Supplementary Material) in order to obtain equilibrated DNA conformations at varied target elongations - namely, *R*_*ee*_ = 1000, 1100, 1200 nm, with *R*_*ee*_ the DNA end-to-end distance. These conformations are 76%, 84% and 92% of the contour length *L* (≈ 1310 nm) of the chain. Indeed, the desired (average) elongation ratios were maintained by applying a harmonic potential to one end of the DNA molecule, with harmonic constant *k* = 5.7 N/m, and an external force *F*_*z*_ = 0.42, 0.88, 3.06 pN to the terminal nucleotides at the opposite end, derived as follows [38]:

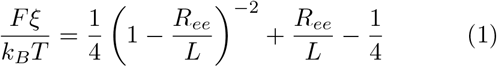

with ξ ≈ 50 nm the persistence length of DNA. Such conformations are compliant with a low-force regime (≤ 5 pN), i.e. where DNA reacts by entropic compliance to the applied tension and the Marko-Siggia formulation of the worm-like chain model is applicable [31, 39]. The three configurations of the DNA helix served as starting frames of a set of classic MD replicas aimed at assessing the early effects of DSBs.

As the radiation-enforced lesions of DNA bear several levels of complexity varying upon the resolution scale [3, 16], we adopt a simplified, effective framework to characterize the local strain from the disruption of the DNA backbone that both faithfully depicts the process and is numerically feasible. Indeed, we focused on T1-DSBs as described by Schipler and Iliakis [3], where the phosphoester bonds of the DNA backbone are disrupted. We thus neglect all modifications involving the nitrogenous bases, and allow thermal fluctuations to detach the DNA moieties.

We hereby define the DSB distance *b*_*d*_ as the distance in-between of the twofold nicks of the DNA backbone, quantified in number of native base pairs (Fig. 1). Diverse DSB *motifs* were enforced by removing the covalent bonds between adjacent nucleotides on both DNA strands at distances *b*_*d*_ = 0, 1, 2 and 3, about halfway of the DNA filament (see Section **1** in the Supplementary Material): in LAMMPS, this is implemented by deleting the bond ID between nearest neighbouring nucleotides in the topology file. For each value of *b*_*d*_, numerous independent replicas were run from the same starting configurations. We remark that no prior minimisation/thermalization protocol of the DSB scenarios was carried out, to depict the subsequent dynamics of a DNA molecule broken abruptly by the radiation field. A snapshot of a DNA rupture by a DSB at *b*_*d*_ = 1 is shown in Fig. 1.

**FIG. 1.**
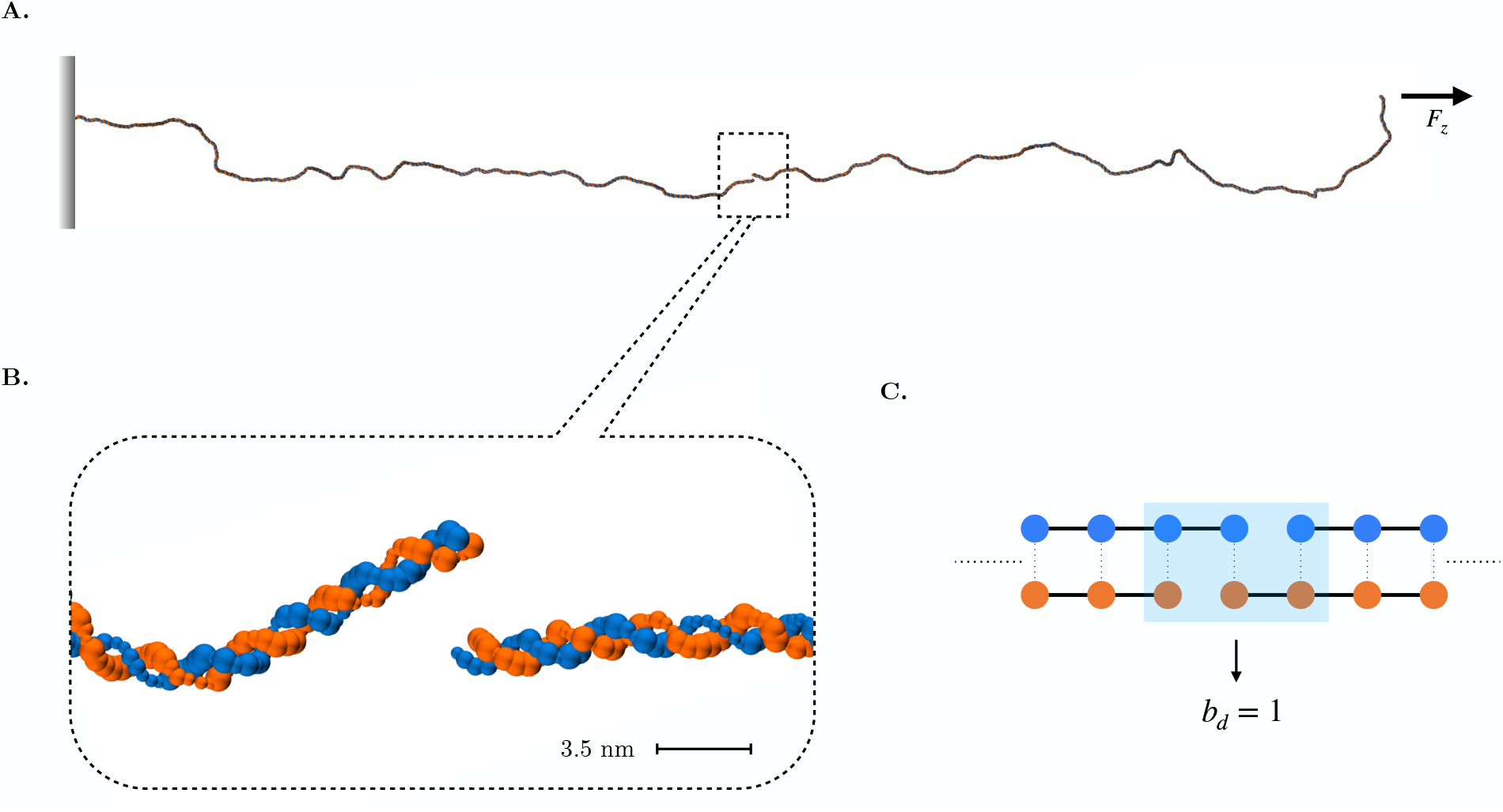
(**A**) Snapshot of the 3855-bp B-DNA molecule associated with an end-to-end distance of ⟨*R*_*ee*_⟩ = 1000 nm, broken by a double-strand break at distance *b*_*d*_ = 1 - shown in detail in the inset (**B**). *F*_*z*_ is the constant external force acting on the DNA, computed according to Eq. 1 -details in the text. (**C**) Schematic depiction of a DNA double-strand break: the backbone connectivity is shown as solid lines, hydrogen bonds between DNA strands as dotted lines. The blue shaded region highlights the nucleotides involved with the residual contact interface between the DNA moieties. To enforce a DSB, covalent bonds are removed between adjacent nucleotides from both DNA strands.

### C. Derivation of the characteristic times of the DNA rupture by DSBs

The characteristic time associated with the rupture of the DNA moieties 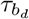 was estimated for all replicas of each (*b*_*d*_, *F*_*z*_) DSB scenario *via* a twofold procedure. We first established a geometrical, heuristic criterion to unequivocally frame the DNA fracturing event, based upon the minimum distance between the broken DNA moieties (see Section **3** Supplementary Material for details): the specific choices were corroborated by visual inspection of the MD trajectories, so that we could focus on a time-frame about the rupturing event. Likewise, we derived 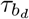 from the internal energy profile of the nucleotides involved with the residual contact interface between DNA moieties at the site of a DSB lesion, that is i) the nucleotides lying in-between the twofold break of the DNA backbone and ii) the nucleotides closely overlooking the DSB interface on both sides of the lesion (see Fig. 1C). Fig. 2 shows the internal energy contribution of the contact nucleotides at a blunt DSB (i.e. at *b*_*d*_ = 0), within a time span involving the rupture of the DNA molecule. By inferring that the rupture of DNA by DSBs is arguably a two-state process (as observed by the internal energy profiles), and that the barrier dividing the bound (metastable) and broken thermodynamic basins is symmetrical at the peak, we performed a sigmoidal fit on the internal energy profile as:

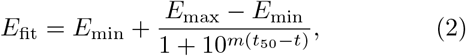

and derived the characteristic time of the DNA rupture at the flex of the curve. Here, *E*_min_ and *E*_max_ are associated with the lower and upper plateau of the internal energy (characterizing the bound and broken states of the DNA molecule, respectively), and *m* is the steepness of the curve. The values of 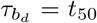 are estimated at the flex of the internal energy transition and closely match those derived by the heuristic criterion, thus validating Δ*U* as an effective proxy to frame the kinetics of the process. As a consequence, we implicitly estimate the internal energy contribution to the process as the difference between the internal energy at the flex of the sigmoid and at the lower energy plateau -which we will tentatively associate with the activation state of the DNA rupture (*vide infra*). Additional and technical details about the fitting protocol are reported in Section **3** of the Supplementary Material.

**FIG. 2.**
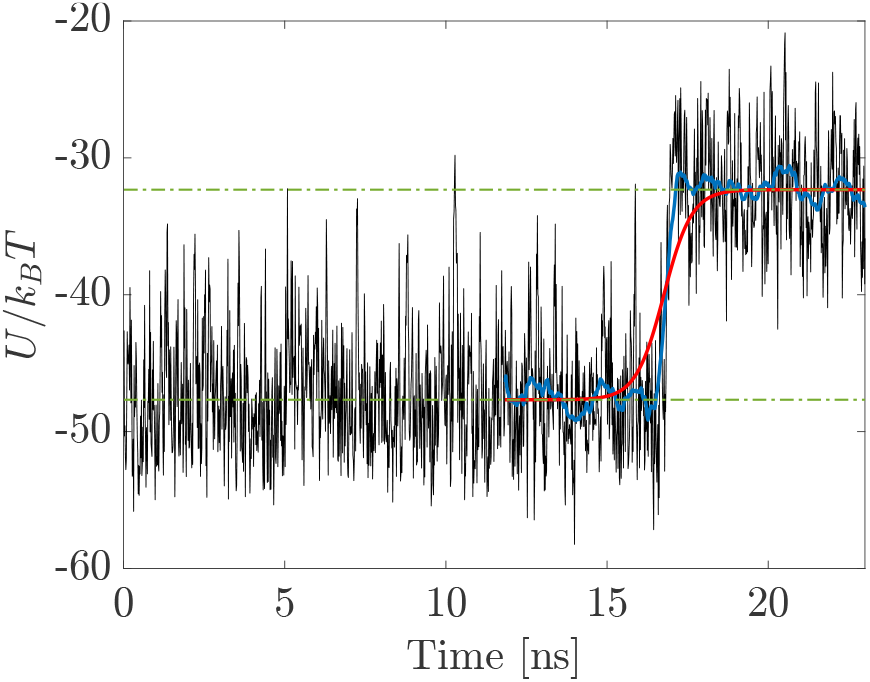
Profile of the local internal energy contribution from the nucleotides involved with the residual contact interface between broken DNA moieties, associated with the DNA rupture by a blunt DSB (*b*_*d*_ = 0) and an average end-to-end distance of the DNA filament ⟨*R*_*ee*_⟩ = 1000 nm. The lesion destabilizes the helical layout, driving a conformational transition of the DNA molecule from a bound to a broken state (lower and upper energy plateaus, shown as dashed green lines). The moving average of the potential energy profile (blue line) is fitted by a sigmoid curve (red line), whereby we assess the characteristic time and an internal energy contribution to the DNA rupture.

## III. RESULTS AND DISCUSSION

As the structure of the DNA double helix is compromised by a DSB lesion, non-covalent interactions hold the broken DNA moieties: the system is thus confined to a metastable, bound state, until the kinetic barrier associated with the rupture of the residual contact interface between the DNA moieties is overcome by means of stochastic thermal fluctuations.

### A. Kinetics of the DNA fracture by DSB motifs

We firstly assessed the kinetics of the DNA fracture from the diverse DSB motifs by estimating the average rupture times 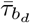 over all independent MD replicas associated with a given (*b*_*d*_, *F*_*z*_) combination -results are outlined in Table I.

**TABLE I.** **Left:** Internal energy contribution to the DNA rupture as function of the DSB distance *b*_*d*_ and applied force, the latter being associated with an average end-to-end distance of the DNA molecule ⟨*R*_*ee*_⟩. **Right:** Average rupture times 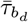 from the different DSB motifs and DNA extensions: each value of 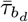 is estimated by averaging over all independent MD replicas. The characteristic times are scaled according to a straightforward conversion of LJ into SI units [34]. In all cases, standard deviations are on the order of the average values due to the statistical nature of the process (see Supplementary Materials **5**).

| $F_z$ [pN] | $\langle R_{ee} \rangle$ [nm] | $\Delta\bar{U}/k_B T$ | | | | $\bar{\tau}$ [ns] | | | |
| --- | --- | --- | --- | --- | --- | --- | --- | --- | --- |
| | | $b_d = 0$ | $b_d = 1$ | $b_d = 2$ | $b_d = 3$ | $b_d = 0$ | $b_d = 1$ | $b_d = 2$ | $b_d = 3$ |
| 0.42 | 1000 | $8 \pm 2$ | $14 \pm 4$ | $23 \pm 4$ | $27 \pm 6$ | $1 \pm 1$ | $6 \pm 6$ | $105 \pm 102$ | $139 \pm 123$ |
| 0.88 | 1100 | $7 \pm 2$ | $14 \pm 4$ | $24 \pm 4$ | $28 \pm 6$ | $0.8 \pm 0.7$ | $4 \pm 4$ | $105 \pm 104$ | $138 \pm 120$ |
| 3.06 | 1200 | $8 \pm 2$ | $14 \pm 4$ | $24 \pm 4$ | $27 \pm 4$ | $0.9 \pm 0.9$ | $6 \pm 7$ | $88 \pm 89$ | $114 \pm 102$ |

The estimates of the average rupture times 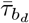 seemingly fit an exponential law of the DSB distance - or equivalently, by a linear function of *b*_*d*_ as:

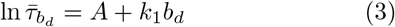

with *A* and *k*_1_ coefficients from the fitting procedure - reported in Table II and shown in Fig. 3. This trend replicates the exponential increase with size observed by Sengar and co-workers for the dissociation kinetics of short DNA duplexes, employing an earlier version of the oxDNA force field [34], thereby characterizing the rupture of DNA by DSBs as a rare event. Notably, our data are similarly well-fit by an alternative law of the form 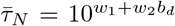 (see Section **6** in the Supplementary Material), which has been reported for the apparent thermal off-rates from the forced melting of DNA duplexes by AFM spectroscopy, extrapolated in the limit of zero force [40–42].

**TABLE II.** *A* 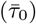 and *k*_1_ (*β*ΔΔ*G*^‡^) coefficients from the fitting of the average rupture time 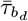 on the DSB distances *b*_*d*_ (Eqs. 3, 4) for each (*b*_*d*_, *F*_*z*_) DNA scenario.

| $F_z$ [pN] | $A$ ($\bar{\tau}_0$) [ns] | $k_1$ ($\beta\Delta\Delta G^\ddagger$) |
| --- | --- | --- |
| 0.42 | $1.0 \pm 1.0$ | $1.7 \pm 0.4$ |
| 0.88 | $0.9 \pm 0.7$ | $1.8 \pm 0.4$ |
| 3.06 | $1.0 \pm 1.0$ | $1.7 \pm 0.4$ |

**FIG. 3.**
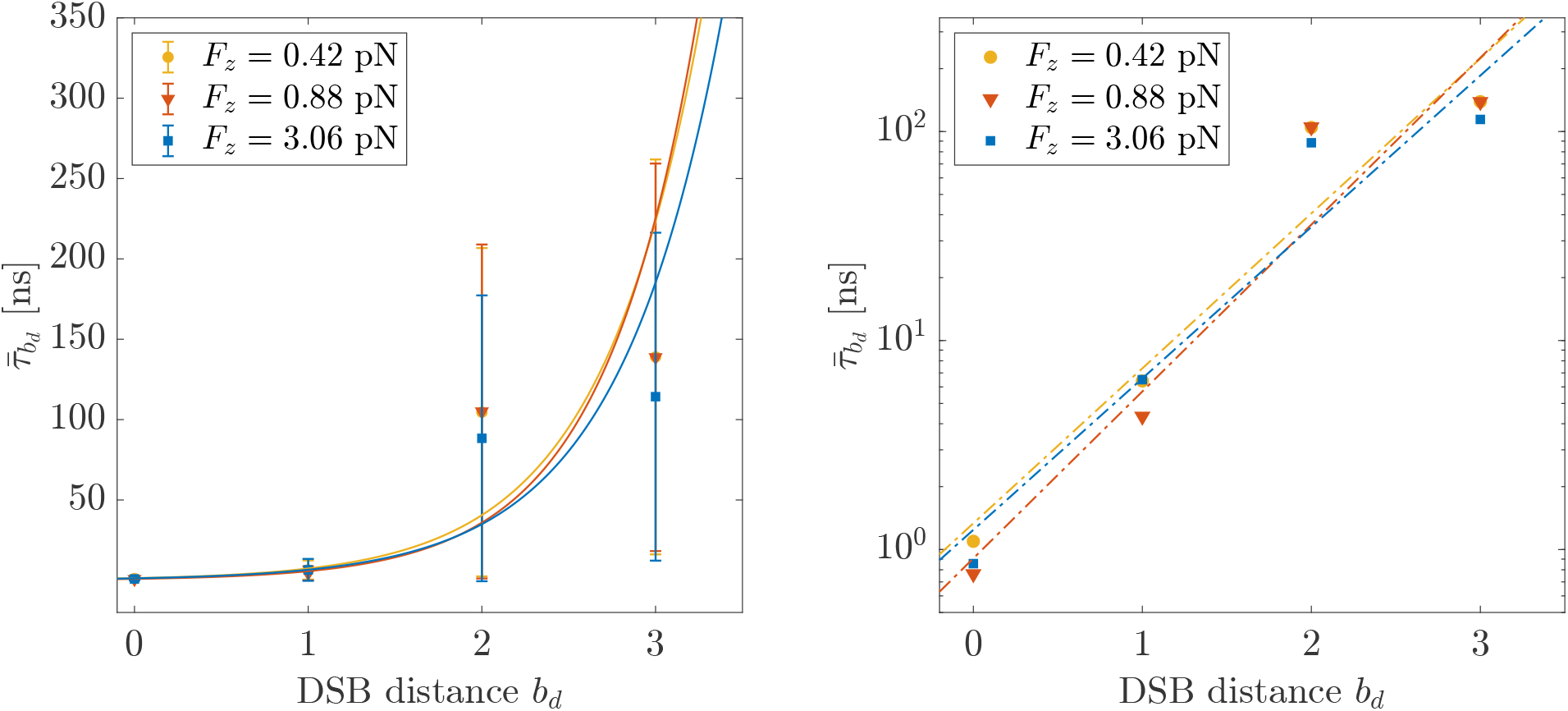
Average rupture times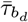 as function of the DSB distance *b*_*d*_ (shown in semi-logarithmic scale in the right panel). The solid and dashed lines are defined upon the fitting protocol described by Eq. 3.

In fact, by assuming the rupture of DNA by DSBs being a cooperative, thermally-activated process (as similarly described in Ref. [41, 42] for the denaturation of DNA), it is tempting to redefine the (*A, k*_1_) coefficients of the exponential law (Eq. 3) in terms of Arrhenius-like factors as:

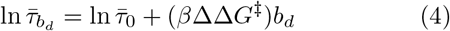

with 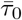 the average fracture time of DNA by blunt DSBs (i.e. at distance *b*_*d*_ = 0), and in a broader sense, the kinetic contribution from the rupture of (stacking, electrostatic) interactions to the activation barrier; *β*^−1^ = *k*_*B*_*T* = 0.62 kcal/mol at 310 K; ΔΔ*G*^‡^ a sequence-nonspecific, additive contribution from the disruption of Watson-Crick hydrogen bonds to Δ*G*^‡^ (i.e. the activation free energy of the rupturing of DNA by the diverse DSB motifs), the latter thus increasing linearly with *b*_*d*_. As shown in Table II, ΔΔ*G*^‡^ ranges between 1.6 *k*_*B*_*T* to 1.9 *k*_*B*_*T* among all scenarios, which is slightly higher than the values reported in Refs. [40, 42, 43] for the activation free energy contribution from the denaturation of DNA: arguably, this is accounted for by a lower entropic contribution from the local fraying of the DNA termini, the latter being dampened by the residual *π*-stack interactions between the DNA moieties in the current scenario. As expected, 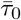 is in line with the estimates of the average rupture times from blunt DSB motifs (see Table I). It might be argued that the application of a constant force regime enhances the kinetic rates of the DNA rupture by lowering the activation free energy barrier along the reaction coordinate of the process [40, 42, 44]. Yet, as shown in Fig. 3 (and further discussed in Section **8** of the Supplementary Material), the average rupture times 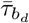 exhibit a negligible dependence upon the tensile protocol enforced on the DNA molecule. In fact, the range of forces applied (and fluctuations thereof) lies within the threshold for a purely entropic response of the DNA molecule under tension [31, 38, 39] and below the mean contribution from the thermal bath (see Section **7.1** of the Supplementary Material). Moreover, the forces exerted at the DNA termini quickly decorrelate along the chain (see Section **7.2** of the Supplementary Material), whereby we infer that thermal fluctuations eventually account for the rupture of the DNA moieties.

### B. Internal energy contribution to the rupture of DNA

We then focused on the internal energy contribution to the rupture of DNA by the DSB motifs and on its dependence on the DSB distance *b*_*d*_. We adopted a consistent criterion (see Section **3** in the Supplementary Material) to define Δ*Ū*(*b*_*d*_) as the average value of the difference between the internal energy of the system at 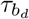 - as described earlier in this work - and the internal energy plateau of the bound basin (values are reported in Table I). Subsequently, we performed a linear fitting as:

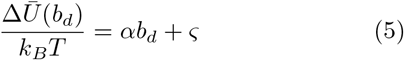

with *α* and *ς* empirical coefficients - reported in Table III. In fact, the validity of this assumption, i.e. the existence of a linear correlation between the internal energy contribution to the rupture of DNA and the DSB distance *b*_*d*_, is verified for all DNA conformations, as shown in Fig. 4. So far, our assessment has thus strongly hinted that the rupture of DNA by double-strand breaks might be depicted effectively by a cooperative, abrupt transition between a metastable, bound state (held by residual hydrogen bonds and stacking contacts) and a broken state of the DNA moieties drifting from the lesion site. Moreover, this is arguably an activated process, well described by transition state theory [45–48], according to which a system progresses between conformational basins across a dynamical bottleneck enforced by the transition state (‡). Given Δ*G*^‡^ as the height of the activation free energy barrier associated with the rupturing process (see Fig. 5), we thus have:

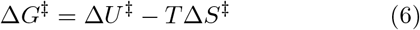

with Δ*U* ^‡^ and Δ*S*^‡^ defining the internal energy and entropic contributions respectively. Taking into account the restraints enforced on the conformational freedom of the DNA molecules, we expect the entropic contribution to be negligible up to the peak of the free energy barrier, thus reducing Eq. 6 to:

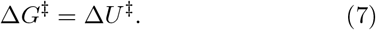

As the barrier is overcome, the DNA moieties acquire a significant conformational freedom, thereby abruptly increasing the entropy of the system -Fig. 5 schematically illustrates the process.

**TABLE III.** *α* and *ς* coefficients from the linear fitting of the average internal energy contribution to the rupture of DNA Δ*Ū*(*b*_*d*_) upon the DSB distance, as described in Eq. 5, at different values of the external force.

| $F_z$ [pN] | $\alpha$ | $\varsigma$ |
| --- | --- | --- |
| 0.42 | $7.1 \pm 1.5$ | $7.8 \pm 1.6$ |
| 0.88 | $7.5 \pm 1.6$ | $7.0 \pm 1.9$ |
| 3.06 | $6.7 \pm 1.4$ | $7.9 \pm 1.7$ |

**FIG. 4.**
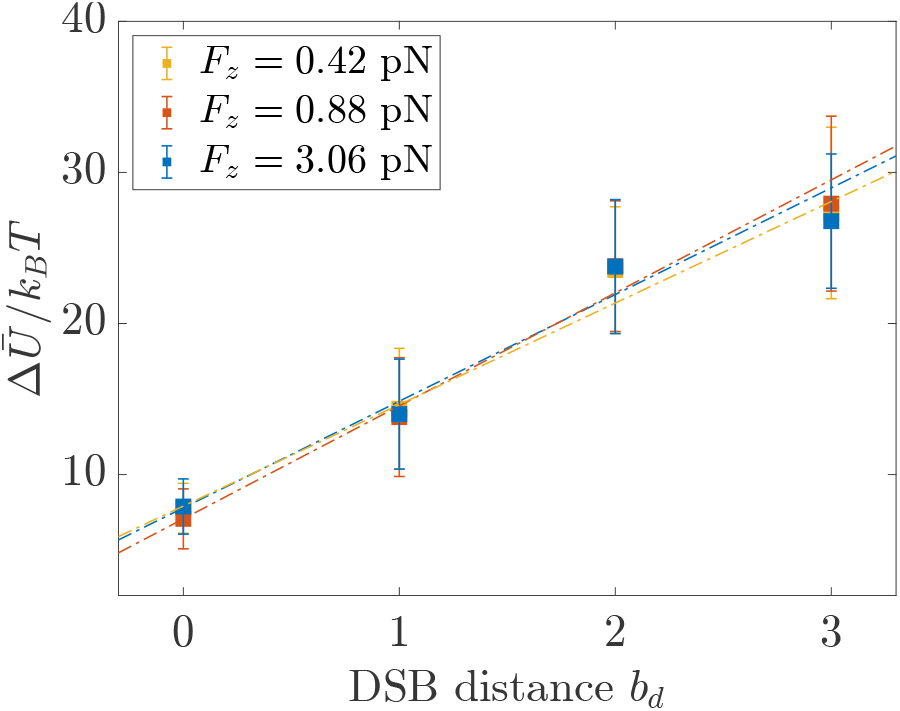
Average internal energy contribution to the rupture of DNA as function of the DSB distance *b*_*d*_ for each force regime. The dashed curves are obtained from the fitting protocol described by Eq. 5.

**FIG. 5.**
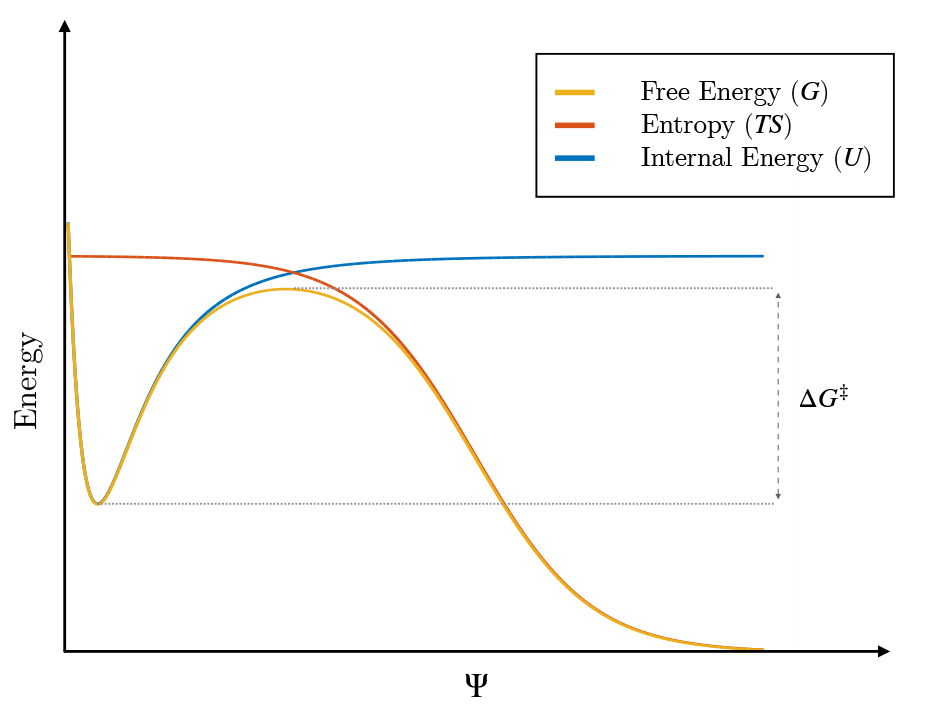
Schematic depiction of the free energy profile (yellow curve) associated with the rupturing process of DNA, as a function of an effective reaction coordinate Ψ. In spite of the twofold nick of the DNA backbone, non-covalent interactions hold the DNA moieties within a locally metastable thermodynamic basin. Subject to a tensile force and limited fraying of the DNA overhangs at the contact interface between the broken moieties, the local conformational freedom of DNA is restrained. A major contribution to the activation free energy of the process is thus provided by the internal energy *U* (blue curve), whereas the entropic contribution *T S* is approximately constant (orange curve). This assumption holds up to the transition state of the rupturing process, described by the activation energy Δ*G*^‡^: In fact, as the DNA moieties drift from the lesion site, several degenerate conformations become available, associated with a significant increase in the entropy of the system.

In light of these assumptions, we tentatively simplify and redefine Δ*U* ^‡^ by the internal energy contribution to the DNA rupturing process. Yet, Δ*Ū*(*b*_*d*_) hardly applies as an effective proxy of the activation free energy: in fact, the ratio between *α* and ΔΔ*G*^‡^ (both representing an additive contribution to the activation energy - see Eqs. 5 and 4 respectively), is about 5-fold. Arguably, this is accounted for by a misestimate of the activation barrier, likely lying closer to the plateau of the metastable, bound state than to the flex of the sigmoidal fit of Δ*U* (*b*_*d*_). Indeed, the rupture of DNA is driven by an abrupt transition, so that small uncertainties in the estimate of 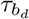 reflect wide variations in the internal energy contribution (see Fig. 2). Moreover, the assumption of a negligible entropic contribution, which is valid up to the peak of the free energy barrier, requires an independent validation through refined MD techniques aimed at characterizing the reaction coordinate and the transition state - which is the objective of ongoing work.

### C. Correlation between physical and simulation time in CG models

It is well acknowledged that CG models enhance the dynamics of systems with respect to their all-atom counterparts [24, 49, 50]. This is inherent to the integration of several degrees of freedom, whereby the coarse-graining procedure somewhat smoothens the free energy landscape, thus favoring the transition between local free energy minima. Furthermore, CG models (such as oxDNA) are frequently implemented with implicit solvents, which lack a proper characterization of hydrodynamic effects from the collective motion of the solute particles nearby [51, 52]. These limitations only marginally, if at all, affect the characterization of the structural features of a system at equilibrium [31, 53]; they become critical, however, to the dynamical assessment of kinetic observables. Under the assumption that the modifications induced by the coarse-graining protocol are quantifiable in terms of a global redefinition of the characteristic times of a system [49], these issues might be tackled by rescaling the time steps of the CG simulation to match the outcome from a reference scenario, such as all-atom MD or wet lab experiments.

In this view, we related the timescales of the physical (experimental) process and those of its *in silico* (CG) depiction *via* the ratio of diffusion coefficients of double-stranded DNA molecules Γ(*ζ*) as:

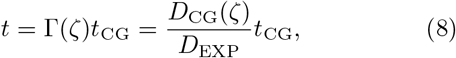

where *t* is the best estimate of the time of the physical process, *t*_CG_ is the time of the CG simulation, and *D*_CG_(*ζ*) and *D*_EXP_ are the coarse-grained and experimental diffusion coefficients of DNA respectively. In fact, a rescaling protocol based on the diffusion coefficients of DNA was adopted by Ouldridge and co-workers alike, who explored the hybridization kinetics of DNA oligomers [54].

To estimate the value of *D*_CG_(*ζ*) for the freely-diffusing 3855-bp DNA molecule, we performed a MD simulation of 5*×*10^6^ *τ* in the NVT ensemble at *T* = 310 K, starting from the equilibrated, 1000-nm conformation of DNA. The value of *D*_CG_ = 2.2 *×*10^−11^ *μ*m^2^*/τ* was thus inferred from a linear fitting of the mean-squared displacement versus time (see Section **4** in the Supplementary Material).

As for *D*_EXP_, we relied on the experimental procedure of Robertson and co-workers [55], who estimated the diffusion coefficient of fragments of *λ*-phage DNA obtained by coupling the action of restriction enzymes and gel electrophoresis. We thus extrapolated the value of *D*_EXP_ from the experimental diffusion coefficients of longer DNA filaments, and scaled it by the ratio of the temperatures adopted in our simulations and in the experimental setup, achieving *D*_EXP_ = 1.17 *μ*m^2^/s and a time-scaling factor Γ(*ζ*) of 11.1.

Results of the rescaling protocol applied to the characteristic rupture times of the DNA molecule at ⟨*R*_*ee*_⟩ = 1000 nm are shown in Table IV, where we artificially extended the application range of Eq. 4, extrapolating the properties of DSB motifs at larger distances (*b*_*d*_ ≥ 4) in a purely speculative effort. By implying that the rupture of DNA is a two-step process regardless of the DSB motif, driven by the abrupt disruption of the residual contact interface between the DNA moieties, we infer a wide kinetic span (10^−8^ -10^−1^ s) associated with DSB distances *b*_*d*_ = 0 10. Despite the limitations and lack of validation of the protocol (and several approximations thereof), our estimates are in fair agreement with the characteristic dissociation times of free DNA duplexes in solution [40] -the latter being likely characterized, however, by a non-negligible entropic contribution from the DNA fraying. In fact, a unique, earlier all-atom MD assessments on the thermal stability of DSBs over short DNA molecules reported of significantly lower kinetic rates, associating the rupture of DNA by DSB motifs at *b*_*d*_ = 3 to characteristic timescales in the order of hours [22]. Such discrepancy might arise from a misestimate of the strength of non-covalent interactions in principle by both (CG, atomistic) models, however knowingly affecting classic atomistic force fields [23, 24, 56].

**TABLE IV.**
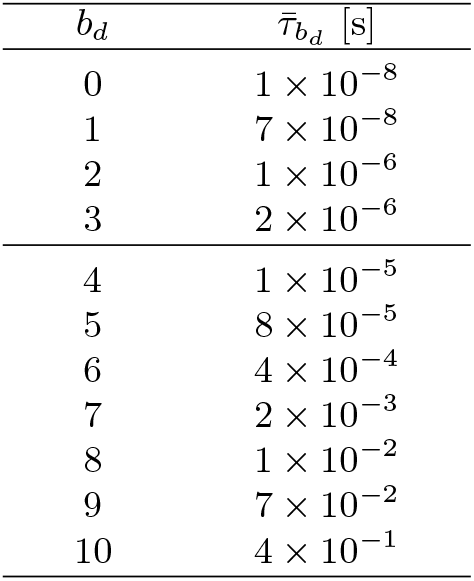
Characteristic times of the DNA rupturing associated with an external force *F*_*z*_ = 0.42 pN, rescaled *via* Eq. 8. Values below *b*_*d*_ = 4 are the (scaled) estimates shown in Table I, whereas the characteristic rupture times relative to higher DSB distances are extrapolated employing Eq. 4

| $b_d$ | $\bar{\tau}_{b_d}$ [s] |
| --- | --- |
| 0 | $1 \times 10^{-8}$ |
| 1 | $7 \times 10^{-8}$ |
| 2 | $1 \times 10^{-6}$ |
| 3 | $2 \times 10^{-6}$ |
| 4 | $1 \times 10^{-5}$ |
| 5 | $8 \times 10^{-5}$ |
| 6 | $4 \times 10^{-4}$ |
| 7 | $2 \times 10^{-3}$ |
| 8 | $1 \times 10^{-2}$ |
| 9 | $7 \times 10^{-2}$ |
| 10 | $4 \times 10^{-1}$ |

## IV. CONCLUSIONS

Double-strand breaks (DSBs), i.e. the twofold cut of the DNA backbone on complementary strands, are a likely and detrimental outcome of the radiation field in cell nuclei. A dynamical characterization of the local breakdown of DNA is thus critical, as it sets the stage for the subsequent cascade of events, involving the activation of the DDR machinery and the rejoining of the broken DNA moieties.

In this work, we tackled the *in silico* assessment of the early stages of the mechanical rupture of DNA by DSBs enforced on a constrained DNA filament. We performed extensive molecular dynamics simulations employing a coarse-grained model of DNA, which let us cover biologically significant timescales, yet keeping a proper level of detail of the molecular process. Moreover, we established a consistent protocol to estimate both the timescales and a characteristic internal energy contribution to the rupturing process from the diverse DSB motifs.

In all scenarios, where the DNA molecule has been kept at varied (average) end-to-end distances by the application of a constant force regime, we observed an exponential increase of the average rupture times with the DSB distance, the fitting coefficients varying negligibly upon the tensile protocol enforced. We thus inferred the DNA rupture by DSBs being a thermally-activated process, as corroborated by the characteristic times being distributed according to an exponential probability density (see Section **5** in the Supplementary Material), which is associated to the likelihood of rare events.

## SUPPLEMENTARY MATERIAL

Supplementary Material is available. Moreover, the raw data associated with this work are freely available on a Zenodo repository at the link https://doi.org/10.5281/zenodo.7941733.

## ACKNOWLEDGMENTS

The authors wish to thank Roberto Menichetti and Luca Tubiana for a critical reading and insightful comments. This project received funding from the European Research Council (ERC) under the European Union’s Horizon 2020 research and innovation program (Grant 758588). RP, LP and MM acknowledge support from the Italian Ministry of Education, University and Research (MIUR) through the FARE grant for the project HAMMOCK (Grant R18ZHWY3NC).

