## Supplemental material for "Kinetics of radiation-induced DNA double-strand breaks through coarse-grained simulations"

#### 1 DNA sequence template

Here, we report the DNA sequence template employed to construct the double-stranded DNA molecule, while Fig. 1 shows the local nucleotide sequence associated with each DSB motif.

```
   1 ATGGTTTATT CTTATACTGA AAAAAAACGT ATTCGTAAAG ATTTTGGTAA ACGTCCTCAA GTTTTAGATG
  71 TTCCTTATTT ATTATCTATT CAATTAGATT CTTTTCAAAA ATTTATTGAA CAAGATCCTG AAGGTCAATA
 141 TGGTTTAGAA TTTCGTTCTG TTTTTCCTAT TCAATCCTAT TCCGGTAATT CCGAATTGCA ATATGTTTCC
 211 TATCGCTTGG GTGAACCTGT TTTTGATGTT CAAGAATGTC AAATTCGCGG TGTTACTTAT TCCCCTTTGC
 281 GCGTTAAATT GCGCTTGGTT ATTTATGAAC GCGAACCTGA AGGTACTGTT AAAGATATTA AAGAACAAGA
 351 AGTTTATATG GGTGAAATTC CTTTAATGAC TGATAATGGT ACTTTTGTTA TTAATGGTAC TGAACGTGTT
 421 ATTGTTTCTC AATTACATCG TTCTCCTGGT GTTTTTTTTG ATTCTGATAA AGGTAAAACT CATTCTTCTG
 491 GTAAAGTTTT ATATAATCGT ATTATTCCTT ATCGTGGTTC TTGGTTAGAT TTTGAATTTG ATCCTAAAGA
 561 TAATTTATTT GTTCGTATTG ATCGTCGTCG TAAATTACCT ACTATTATTT TACGTTTAAA TTATACTACT
 631 GAACAAATTT TAGATTTATT TTTTGAAAAA GTTATTTTTG AAATTCGTGA TAATAAATTA CAAATGGAAT
 701 TGGTCCCCGA ACGCTTGCGC GGCGAAACCT CCTTTGATAT TGAAAATGGC AAAGTCTATG TCGAAAAAGG
 771 CCGCCGCATT ACCCGCCATA TTCGCCAATT GGAAAAAGAT GATGTCAAAT TGATTGAAGT CCCCGTCGAA
 841 TATATTGGCA AAGTCGTCAA AGATTATATT GATGAATCCA CTGGTGAATT GATTTGTAAT ATGGAATTGT
 911 CCTTGGATTT GTTGAAATTG TCCCAATCCG GTCATAAACG CATTGAAACT TTGTTTACTA ATGATTTGGA
 981 TCATGGTCCT TATATTTCCG AAACTTTGCG CGTTGATCCT ACTAATGATC GCTTGTCGCT CGTAGAGATC
1051 TACCGGATGA TGCGGCCAGG AGAGCCACCA ACACGGGAGG AGTCGCTCTT CGAGAACCTC TTCTTCTCGG
1121 AGGACCGGTA CGACCTCTCG GTAGGACGGA TGAAGTTCAA CCGGTCGCTC CTCCGGGAGG AGATCGAGGG
1191 ATCGGGAATC CTCTCCAAAG ATGATATTAT TGATGTTATG AAAAAATTGA TTGATATTCG CAATGGTAAA
1261 GGTGAAGTTG ATGATATTGA TCATTTGGGT AATCGCCGCA TTCGCTCCGT TGGTGAAATG GAAAATCAAT
1331 TTCGCGTTGG TTTGGTTCGC GTTGAACGCG TTAAAGAACG CTTGTCCCTG GGGGACCTGG ACACGCTGAT
1401 GCCGCAGGAC ATGATAAACA AGCCGATAAG CGTGAAGGAG TTCTTCGGGA GCAGCCAGCT GAGCCAGTTC
1471 ATGGACCAGA ACAACCCGCT GAGCGAGATA ACGCACAAGA GGAGGATAAG CCTGGGGCCG GGGGGGCTGA
1541 CGAGGGAGAG GGGGTTCGAG GTGAGGGACG TGCACCCGAC GCACTACGGG AGGGTGTGAC CGATAGAGAC
```

*

1611 GCCGGAGGGG CCGAACATAG GGCTGATAAA CAGCCTGAGC GTGTACCAGA CGAACGAGTA CGGGTTCCTG
1681 GAGACGCCGT ACAGGAAGGT GACGGACGGG GTGGTGACGG ACGAGATACA CTACCTGAGC ATAGAGGAGG
1751 GGAACTACGT GATACAGAAC AGCAACCTGG ACGAGGAGGG GCACTTCGTG GAGGACCTGG TGACGTGAAG
1821 GAGCAAGGGG GAGAGCAGCC TGTTCAGCAG GGACCAGGTG GACTACATGG ACGTGAGCAC GCAGCAGGTG
1891 GTTTCCGTTG GTTCCTTGAT TCCTTTTTTG GAACATGATG ATAATCGCTT GATGGGTAAT ATGCAACGCC
1961 AAGTTCCTAC TTTGCGCGAT AAACCTTTGG TTGGTACTGG TATGGAACGC GTTGTTGATT CCGGTGTTAC
2031 TGTTAAACGC GGTGGTGTTG TTCAATATGT TGATTCCCGC ATTGTTATTA AAGTTAATGA AGATGAAATG
2101 TATCCTGGTG AAGGTATTGA TATTTATAAT TTGACTAAAT ATACTCGCTC CAATCAAAAT ACTTGTATTA
2171 ATCAAATGCC TTGTGTTTCC TTGGGTGAAC CTGTTGAACG CGGTGATGTC TTGGATGGCC CCTCCACCGA
2241 TTTGGGCGAA TTGTTGGGCC AAAATATGCG CGTCTTTATG CCCTGGAATG GCTATAATTT TGAAGATTCC
2311 ATTTTGGTCT CCGAACGCGT CGTCCAAGAA GATCGCTTTA CCACCATTCA TATTCAAGAA TTGTGTGTCT
2381 CCCGTGATAC TAAATTAGGT CCTGAAGAAA TTACTGATAT TCCTAATGTT GGTGAATTAT CTAAATTAGA
2451 TGAATCTGGT ATTGTTTATA TTGGTGAAGT TACTGGTGGT GATATTTTAG TTGGTAAAGT TACTCCTAAA
2521 GGTGAAACTC AATTAACTCC TGAAGAAAAA TTATTACGTA TTTTTGGTGA AAAATCTGAT GTTAAAGATT
2591 CTTCTTTACG TGTTCCTAAT GGTGTTTCTG GTACTGTTAT TGATGTTCAA GTTTTTACTC GTGATGGTGT
2661 TGAAAAAGAT AAACGTTTAG AAATTGAAGA AATGCAATTA AAACAAAAAA AAGATTTATC GGAAGAACTC
2731 CAAATCCTCG AAGGCCTCTT TTCGCGGATC CGGGTCCTCG TCGGCGGCGT CGAAGAAAAA CTCGATAAAC
2801 TCCCCCGGGA TCGGTGGCTC GAACTCGGCC TCACCGATGA AGAAAAACAA AATCAACTCG AACAACTCGA
2871 ACAATATGAT GAACTCAAGC ACGAGTTCGA GAAGAAGCTC GAGAAGCGGC GGAAGATCAC ACAGGGAGAC
2941 GACCTCCCAG GAGTACTCAA GATCGTAAAG GTATACCTCG TAAAGCGGCG GATCCAGCCA GGAGACAAGA
3011 TGGGACGGCA CGGAAACAAG GGAGTAATCT CGAAGATCAA CCCAATCGAG GACATGCCAT ACGACGAGAA
3081 CGGAACACCA GTAGACATCG TACTCAACCC ACTCGGAGTA CCATCGCGGA TGAACATCGG ACAGATCCTC
3151 GAGACACACC TCGGAATGAA GGGAATCGGA GACAAGATCA ACATGCTCAA GCAGCAGCAG GAGGTAAAGC
3221 TAAGAGAGTT CATACAGAGA TACGACCTAG GAGACGTAAG ACAGAAGGTA GACCTAAGTA CATTCAGTGA
3291 CGAGGAGGTA ATGAGACTAG AGAACCTAAG AAAGGGAATG CCAATAACAC CAGTATTCGA CGGAAAGGAG
3361 GAGATAAAGG AGCTACTAAA GCTGGGGGAC CTGCCGACGA GCGGGCAGAT AAGGCTGTAC GACGGGAGGA
3431 CGGGGGAGCA GTTCGAGAGG CCGGTGACGG TGGGGTACAT GTACATGCTG AAGCTGAACC ACCTGGTGGA
3501 CGACAAGATG CACAGGAGCA CGGGGAGCTA CAGCCTGGTG ACGCAGCAGC CGCTGGGGGG TAAACAATTT
3571 GGTGGTCAAC GTTTTGGTGA AATGGAAGTT TGGTTAGAAT ATGGTTATAC TTTACAAGAA ATGTTAACTG
3641 TTAAATCTGA TGATGTTAAT GGTCGTACTA AAATGTATAA AAATATTGTT GATGGTAATC ATCAAATGGA
3711 ACCTGGTATG CCTGAGAGCT TCAACGTGCT GCTGAAGGAG ATAAGGAGCC TGGGGATAAA CATAGAGCTG
3781 GAGGACGAGG AGAGCTTCAA CGTGCTGCTG AAGGAGATAA GGAGCCTGGG GATAAACATA GAGCTGGAGG
3851 ACGAG

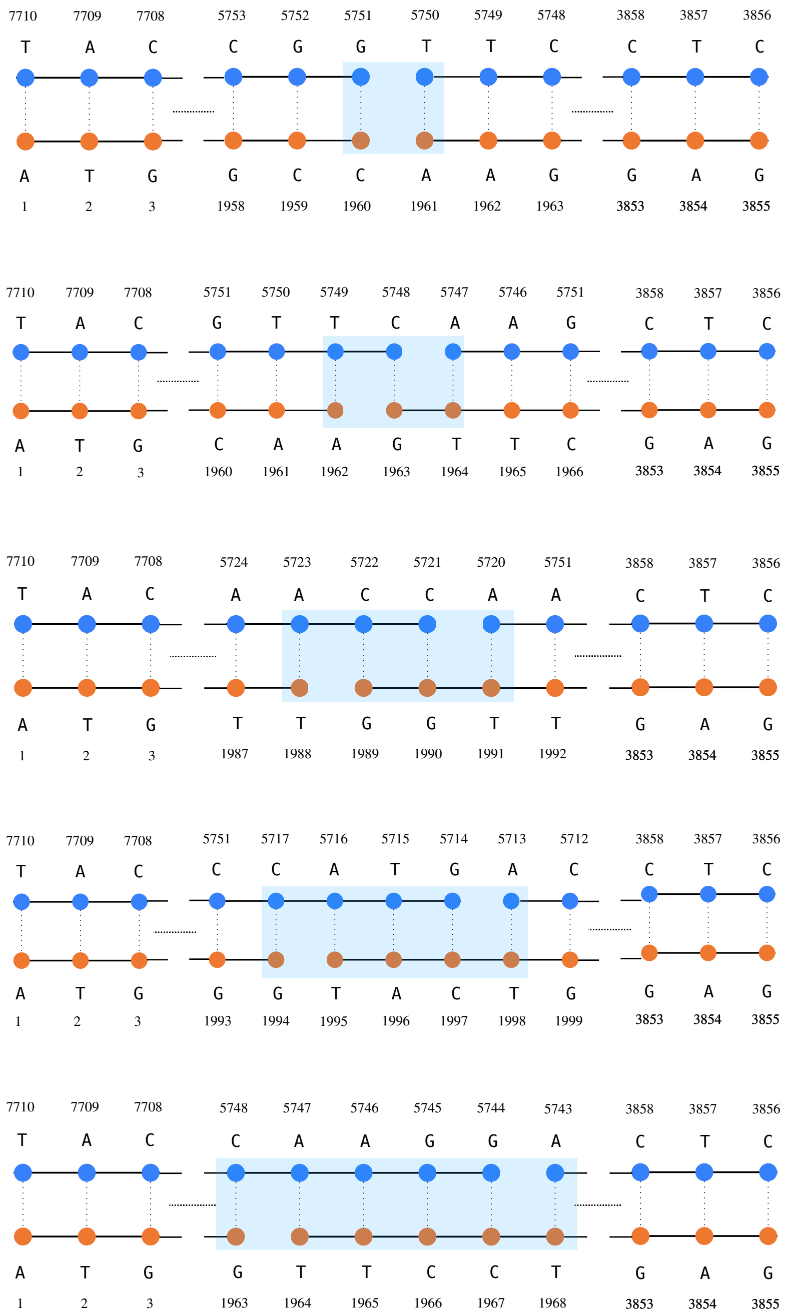


**Supplementary Figure 1:** Schematic depiction of the DSB motifs. Nucleotides are shown as circles; covalent bonds of the DNA backbone as solid lines; Watson-Crick hydrogen bonds between the DNA strands as dotted lines. To enforce a DSB, covalent bonds are removed between adjacent nucleotides from both DNA strands. Each nucleotide is associated with the residue type (A, T, C, G) and the respective LAMMPS atom-ID.

#### 2 Molecular dynamics simulations: DNA thermalization protocol

We carried out a step-wise thermalization protocol as follows: i) we let the DNA molecule relax at $T = 273$ K upon reaching a plateau of the total energy of the system; ii) we heated the system to a working temperature of $T = 310$ K in a quasi-static manner, by performing a single run of $1.6 \times 10^5\ \tau$ in the NVT ensemble, associated with a damping coefficient of the Langevin thermostat $\zeta = 10\ \tau$; iii) we finally ran a subsequent simulation of $3.3 \times 10^5\ \tau$ employing an underdamped Langevin thermostat ($\zeta = 100\ \tau$), to relax the molecule into an equilibrated conformation.
We carried out several instances of steered MD, relaxing the DNA molecule at different elongations (or tensile protocols) - namely, $R_{ee} = 1000,\ 1100,\ 1200$ nm, with $R_{ee}$ the end-to-end distance of the DNA chain [6].
Such conformations were obtained by employing soft harmonic potentials restraining both DNA termini, associated with elastic constants $k_1 = 5.7$ N/m and $k_2 = 2.3 \times 10^{-3}$ N/m respectively, at $\zeta = 10\ \tau$. This protocol allowed us to slowly stretch the thermalized DNA molecule to each of the target elongations. Upon reaching the desired DNA conformation, we thus performed subsequent equilibration stages of i) $4.9 \times 10^4\ \tau$ at $\zeta = 100\ \tau$, and ii) $5.8 \times 10^5\ \tau$ at $\zeta = 10\ \tau$. The three configurations of the DNA helix served as initial frames of a set of classic MD runs aimed at assessing the early non-equilibrium effects of DSBs.

### 3 Heuristic assessment of the rupture of DNA by a DSB

Double-strand breaks (DSBs) are enforced on a DNA molecule by removing the covalent bonds between adjacent nucleotides over complementary strands - that is, between $(n_i, n_{i+1})$ and between $(n_j, n_{j+1})$. To devise a consistent criterion that unequivocally characterizes the rupture of DNA, we ascertain the disruption of all residual contacts between the DNA moieties by satisfying the following relation:

$$\min\{r_{i,i+N+1}, r_{j,j+N+1}\} > 2 \times b_d \langle r_{\mathrm{B}} \rangle + \Lambda, \tag{1}$$

where $r_{i,i+N+1}$ and $r_{j,j+N+1}$ are distances between indexed nucleotides (shown in Fig. 2), and $\langle r_{\mathrm{B}} \rangle$ is the equilibrium distance between adjacent nucleotides from a $2.5 \times 10^6$ $\tau$, NVT simulation of the intact 3855-bp DNA molecule held at an end-to-end distance of 1000 nm. $\Lambda$ is defined heuristically *via* a recursive protocol accounting for all MD trajectories, thus fulfilling a consistent criterion across all DSB motifs (as shown in Fig. 3), and has been set to 1.7 nm.

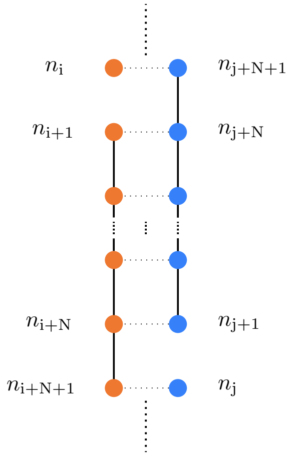


**Supplementary Figure 2:** Schematic depiction of a DSB motif at an arbitrary distance $b_d$ = N. Nucleotides are shown as circles; covalent bonds of the DNA backbone as solid lines; Watson-Crick hydrogen bonds between the DNA strands as dotted lines. To enforce a DSB, covalent bonds are removed between adjacent nucleotides from both DNA strands (between $(n_i, n_{i+1})$ and $(n_j, n_{j+1})$).

#### 3.1 The fitting protocol

To assess the characteristic times of the DNA rupturing process $\tau_{b_d}$ from all independent simulations, we fitted the data from the variation of internal energy at the residual contact interface between the broken DNA moieties by a step-wise protocol as follows:

- we first employed Eq. 1 to approximately estimate $\tilde{\tau}_{b_d}$, thereby defining a time interval $[t_{\min}, t_{\text{end}}]$, such that $\tilde{\tau}_{b_d} \in [t_{\min}, t_{\text{end}}]$. The range is set to optimize the time frame to fit the internal energy profile;

- to further improve the quality of the fitting, we refined the dataset by performing a moving average of the internal energy contribution within the $[t_{\min}, t_{\text{end}}]$ frame;

- we thus exploited the **sigm_fit** modulus of Matlab® to fit the internal energy profile $U/k_BT$ by a sigmoidal function of the form:

$$E_{\text{fit}} = E_{\min} + \frac{E_{\max} - E_{\min}}{1 + 10^{m(t_{50}-t)}}, \tag{2}$$

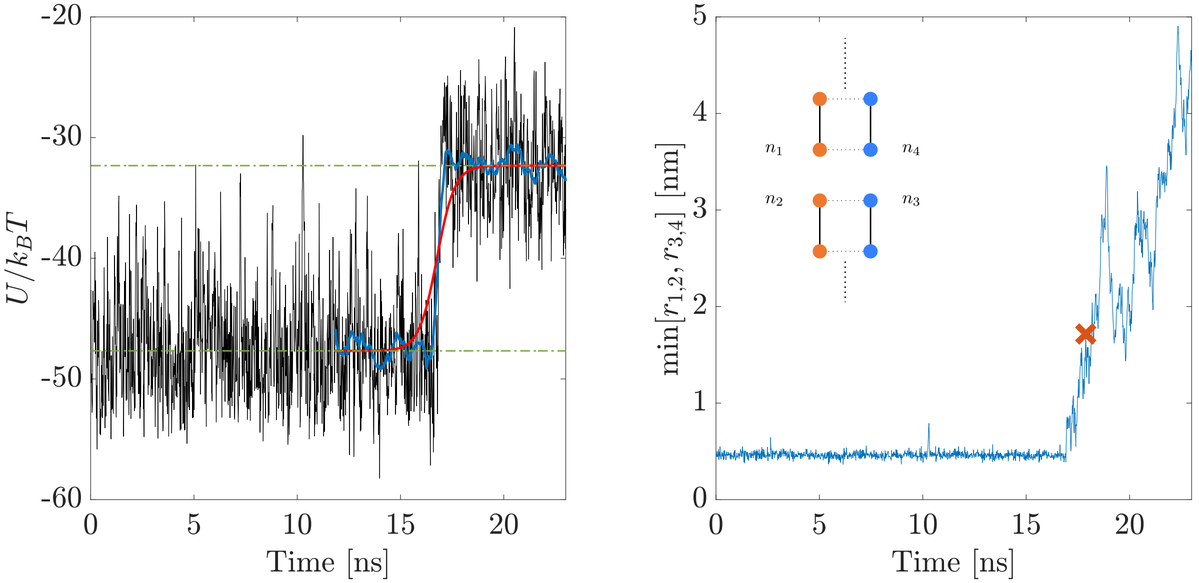


**Supplementary Figure 3:** **Left:** The internal energy contribution associated with the rupture of DNA from a DSB motif at distance $b_d = 0$, enforced on a 3855-bp DNA molecule experiencing an external force $F_z = 0.42$ pN. The moving average of the internal energy (shown in blue) is fitted by a sigmoid curve (Eq. 2 - red line). Dashed green lines highlight the internal energy plateaus of the (metastable) bound and broken states of the DNA molecule. **Right:** Time evolution of the heuristic distance criterion, as described by Eq. 1, associated with the rupture of DNA from a blunt DSB motif, i.e. at distance $b_d = 0$ (details in the text) - the threshold set by Eq. 1 is highlighted by a red mark.

where $E_{\text{min}}$ and $E_{\text{max}}$ are associated with the lower and upper plateau of the internal energy respectively, $m$ defines the steepness of the curve, and $t_{50} = \tau_{b_d}$ is estimated at the flex of the internal energy transition.

For each DSB scenario, this protocol thus provides us with a characteristic time of the DNA rupturing process $\tau_{b_d}$ (or $t_{50}$), and an estimate of the internal energy of the (metastable) bound and broken states as $E_{\text{min}}$ and $E_{\text{max}}$ respectively. Moreover, we consistently obtain an internal energy contribution to the rupturing process from the sharp transition of $U/k_BT$ as:

$$\Delta U/k_BT = E_{\text{fit}}(\tau_{b_d}) - E_{\text{min}}. \tag{3}$$

Further details on the scripts are available at the Zenodo repository. Lastly, we remark that the estimates of the rupture times from the heuristic criterion (Eq. 1) match those extrapolated from the sigmoidal fitting procedure, thereby validating the fitting protocol.

### 4 Derivation of the CG diffusion coefficient of freely-diffusing DNA

To extract the diffusion coefficient of the 3855-bp DNA molecule in the CG model, we started from a thermalized conformation of the 1000-nm steered DNA molecule and allowed it to diffuse freely. We thus performed a MD simulation of $5 \times 10^6$ $\tau$ in the NVT ensemble, employing a damping coefficient of the Langevin thermostat $\zeta = 2.5$ $\tau$, thereby extracting $D_{\mathrm{CG}}(\zeta)$ from the linear fitting of the the mean-squared displacement (MSD) profile (shown in Fig. 4).

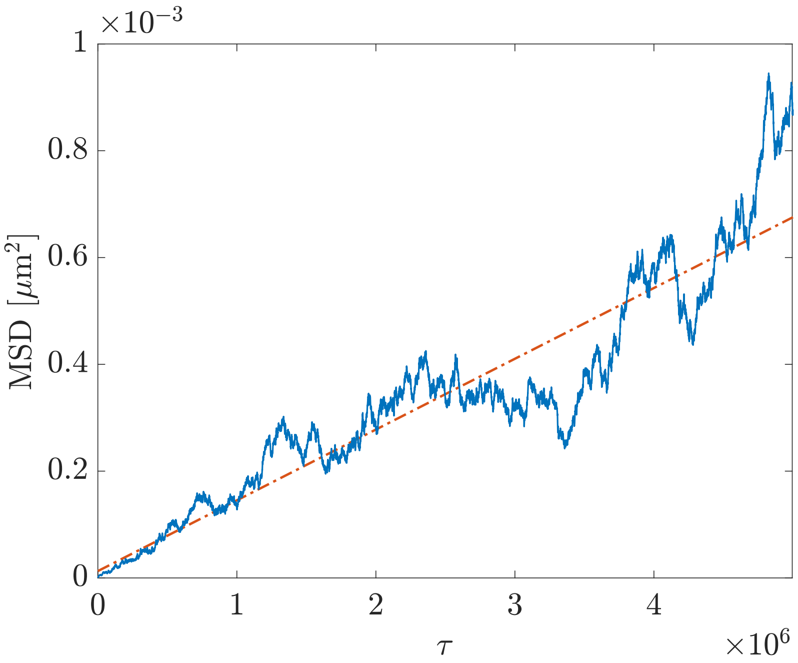


**Supplementary Figure 4:** Linear fitting of the MSD for the freely-diffusing, 3855-bp DNA molecule as a function of time, employing a damping coefficient of the Langevin thermostat $\zeta = 2.5$ $\tau$.

### 5 Statistical analysis of the rupturing events

According to the data shown in Fig. 5, reporting the distribution of the rupturing times for each DSB scenario, we might characterize the rupture of DNA by a DSB as a rare event, associated with a Poisson process. The probability $P_\lambda[t]$ for the DNA molecule to break within an arbitrary time $t$ is defined as:

$$P_\lambda[t] = 1 - \exp\left[-\lambda(b_d)t\right] \tag{4}$$

with $\lambda^{-1}$ the average, characteristic time for the rupturing of DNA by a DSB motif ($b_d$). Thus, the probability density function of the underlying process is exponential.
To verify the reliability of our dataset, we sampled the cumulative distribution function, employing:

$$\frac{N(\tau_{b_d})}{N_0} = 1 - \exp\left[-\lambda\tau_{b_d}\right] \tag{5}$$

with $N(\tau_{b_d})$ the amount of effective simulations detecting the rupture of DNA by time $\tau_{b_d}$, and $N_0$ the total amount of independent MD simulations for a $(b_d, F_z)$ scenario, i.e. either showing a rupture event or hitting wall-time first (maximum threshold of $10^8$ MD steps). Therefore, we performed a fitting procedure over Eq. 5 for each DSB scenario (Fig. 6), thereby extracting the values of $\lambda^{-1}$.

| $F_z$ [pN] | $b_d = 0$ | | $b_d = 1$ | | $b_d = 2$ | | $b_d = 3$ | |
|---|---|---|---|---|---|---|---|---|
| | $\lambda^{-1}$ [ns] | $\langle . \rangle$ [ns] | $\lambda^{-1}$ [ns] | $\langle . \rangle$ [ns] | $\lambda^{-1}$ [ns] | $\langle . \rangle$ [ns] | $\lambda^{-1}$ [ns] | $\langle . \rangle$ [ns] |
| 0.42 | 1 | 1 (488) | 7 | 6 (477) | 103 | 105 (122) | 145 | 139 (106) |
| 0.88 | 0.8 | 0.8 (427) | 4 | 4 (504) | 93 | 105 (102) | 146 | 138 (77) |
| 3.06 | 0.7 | 0.9 (478) | 6 | 6 (525) | 86 | 88 (108) | 119 | 114 (94) |

**Supplementary Table 1:** Comparison between the estimates of the average rupture times from the fitting procedure based on Eq. 5 ($\lambda^{-1}$) and from the MD simulations. The total amount of simulations performed for each scenario is reported in brackets.

As reported in Table 1, the estimates of the characteristic rupture times provided by the fitting procedure match those obtained from the average of the MD simulations.

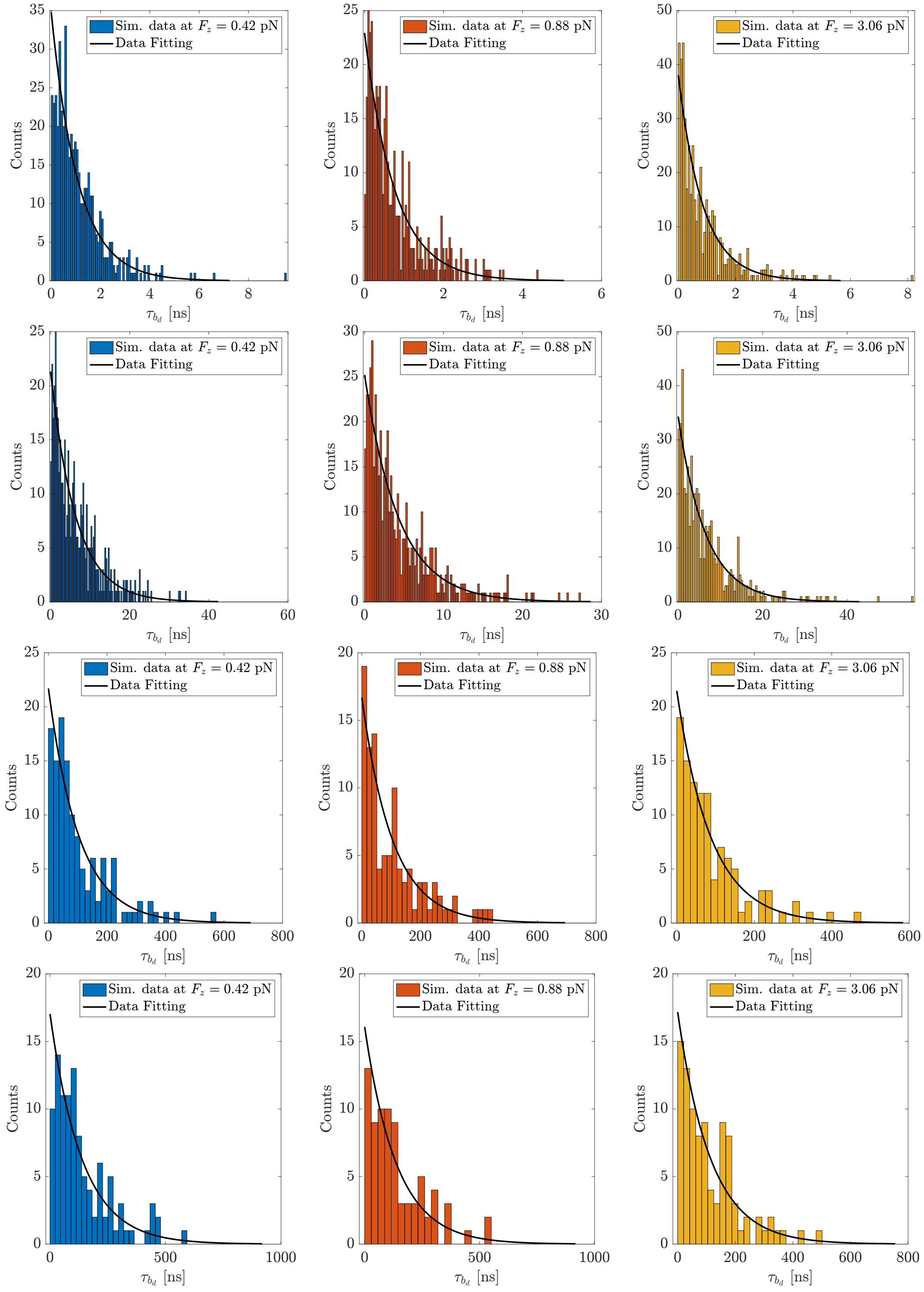


**Supplementary Figure 5:** Histograms of the rupturing times $\tau_{b_d}$, showing the underlying exponential probability density function for different DSB scenarios. Each row is associated with the diverse DSB motifs ($b_d = 0 \div 3$ from top to bottom). The solid black lines highlights the fitting procedure over the corresponding dataset.

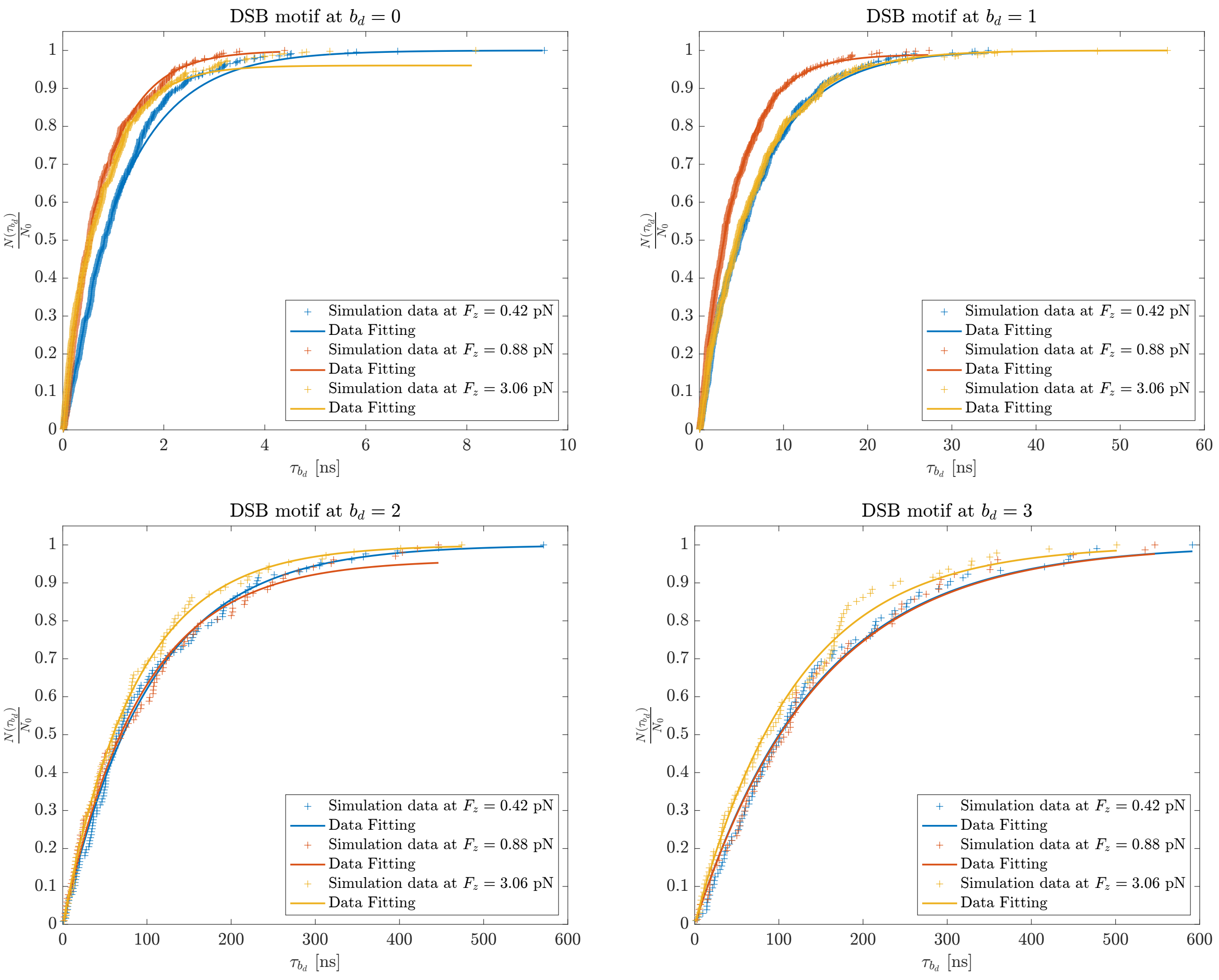


**Supplementary Figure 6:** Cumulative distribution function (as described by Eq. 5) for the different DSB motifs and scenarios - the solid lines show the results from the fitting procedure on the rupture times $\tau_{b_d}$ extracted from the MD simulations.

### 6 Analysis of the forces acting on the system

It is well-acknowledged that the application of an external force generally lowers the free energy barrier along the reaction coordinate of an activated process [2, 3, 4]. However, one might expect that a steering protocol acting in the low-force regime (that is $F_z \leq 5$ pN for the system at hand) should induce conformational changes according to the entropic compliance of the DNA molecule, rather than to an enthalpic, elastic response [1, 6, 5].

To validate the hypothesis that the simulation protocol has a negligible impact on the characteristic rupture times (as well as on the activation energy of the process), thereby allowing the DNA rupturing by thermal fluctuations, we characterized the force profile acting on the DNA molecule.

#### 6.1 Force fluctuations

We performed about 7.5 $\mu$s-long plain MD simulations of intact DNA molecules subject to $F_z = 0.42, 0.88$ and 3.06 pN. For each trajectory, we computed the fluctuations of the total force acting on each DNA centroid, $\sigma^2[|\vec{F}_{tot}|]$, and compared the results against the force contributions from the thermostat alone - the latter defined by the fluctuation-dissipation theorem:

$$\left\langle F_{tot}^2 \right\rangle = \frac{6 k_B T m}{\tau_{in} dt}, \tag{6}$$

with $F_{tot}$ the total force on a DNA centroid, $k_B$ the Boltzmann constant, $T$ the temperature, $m$ the mass of the nucleotide and $\tau_{in}$ the damping coefficient of the thermostat. As shown in Fig. 7, the fluctuations from the two contributions are of the same order of magnitude.

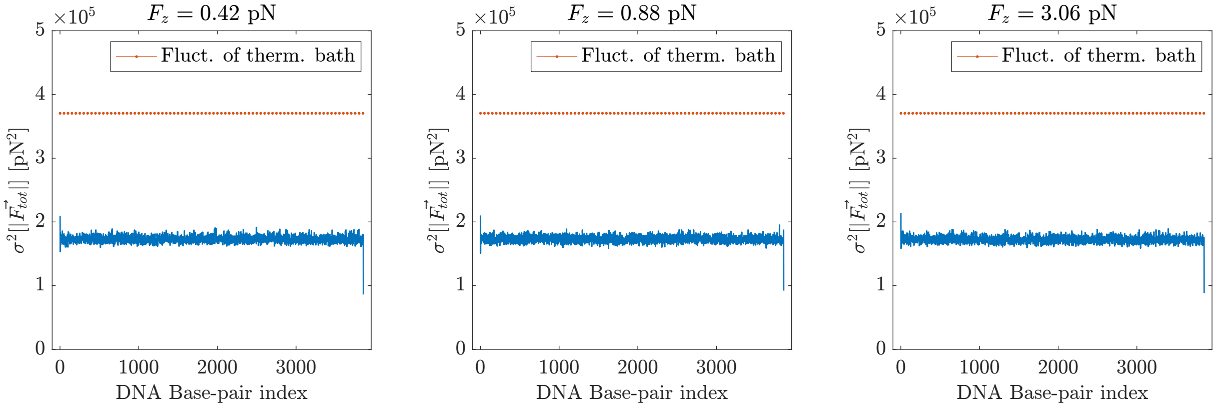


**Supplementary Figure 7:** Fluctuations of the forces acting on each base pair (blue line), against the sole contribution of the Langevin thermostat (orange dashed line), the latter estimated by the fluctuation-dissipation theorem.

#### 6.2 Force correlations along the DNA molecule

We then assessed the propagation of the forces associated with the external force $F_z$ along the DNA molecules, which might eventually enhance the kinetics of the DNA rupture.
Firstly, we performed a 'point-wise' correlation analysis of the forces acting on the centroid of the DNA terminus (i.e. experiencing $F_z$) and on several centroids at various distances along the DNA molecule. The correlation (normalized by the standard deviations of the respective contributions) is defined as:

$$C_{ij}(\tau) = \frac{\langle F_z(i) F_z(j) \rangle}{\sigma[F_z(i)] \sigma[F_z(j)]} = \frac{1}{\sigma[F_z(i)] \sigma[F_z(j)]} \frac{1}{\mathcal{T} - \tau} \sum_{t=0}^{\mathcal{T}-\tau} F_z^i(t) F_z^j(t+\tau), \tag{7}$$

with $i$ and $j$ indexes of the DNA centroids along the chain and $\mathcal{T}$ the total simulation time. Results are depicted in Fig. 8 for the DNA molecule experiencing an external force of $F_z = 3.06$ pN - similar trends are obtained for $F_z = 0.42, 0.88$ pN, where: i) $C_1$ is the auto-correlation of the forces acting on the terminal nucleotides; ii) $C_2$ is the correlation between the forces acting on the terminal nucleotides and the forces acting on nucleotides lying at 150 bp (i.e. the DNA persistence length); iii) $C_3$ is the correlation between the forces acting on the terminal nucleotides and the forces acting on nucleotides lying at 2000 bp (i.e. about the lesion site). Arguably, force correlations are quickly suppressed within a few base pairs.

Subsequently, we employed Eq. 7 to derive the force correlations between DNA centroids at distance $s$ anywhere along the chain:

$$\langle C(\tau) \rangle_s = \frac{1}{N(s)} \sum_s C_{s=|i-j|}(\tau) \tag{8}$$

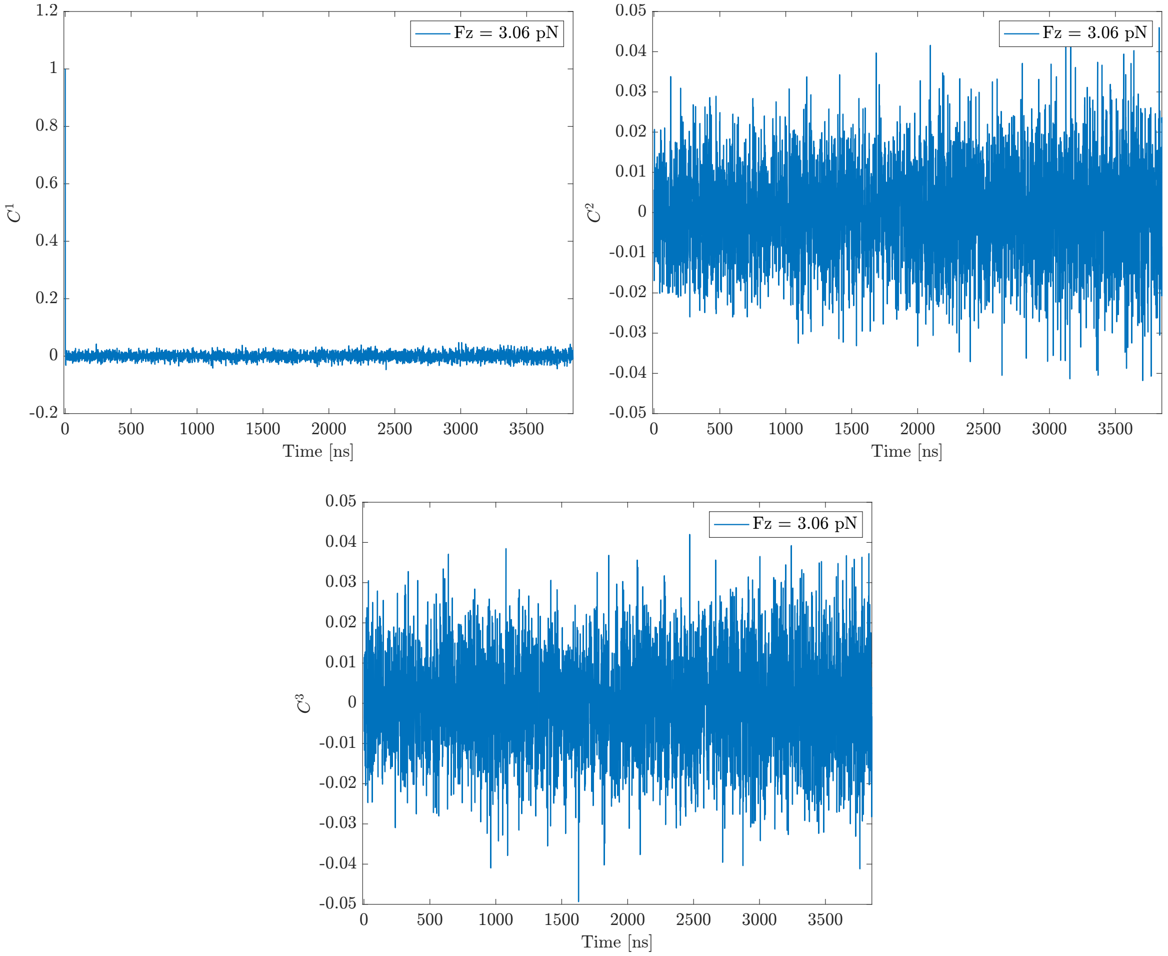


**Supplementary Figure 8: Top-left**: the auto-correlation of the forces acting on the terminal nucleotides (i.e. experiencing an external force of $F_z = 3.06$ pN). **Top-right**, **bottom**: correlation between the forces acting on the terminal nucleotides and on the nucleotides lying at 150 and 2000 bp from the DNA terminus, respectively.

with $N(s)$ the amount of unique $(i, j)$ correlations between nucleotides at distance $s = |i - j|$. Likewise, results are shown in Fig. 9 and report a quickly-decaying correlation of the forces along the DNA molecule.

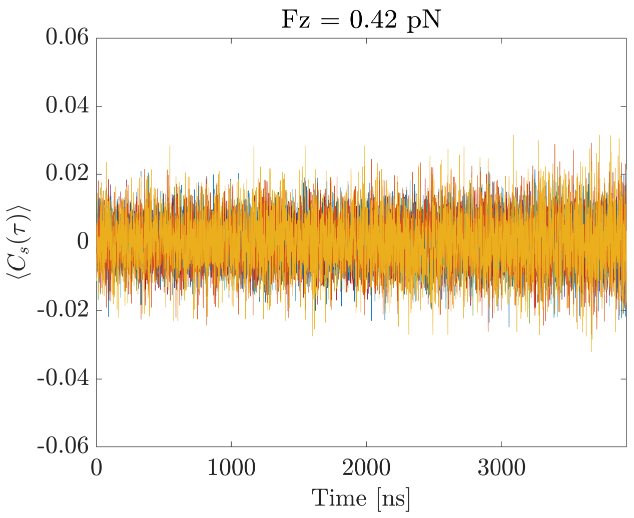


**Supplementary Figure 9:** Average correlations of the forces acting on nucleotides lying at distance $s = 1, 2, ..., 3855$ bp along the DNA chain - tracks are laid on top of one another.

#### 7 DNA rupture kinetics and zero-force extrapolation

To further corroborate the hypothesis that the application of an external force in the low-force regime is associated with a negligible effect on the kinetics of the DNA rupture - so that the process is mostly driven by thermal fluctuations, Fig. 10 shows a (roughly) constant trend of the characteristic rupture times $\bar{\tau}_{b_d}$ as function of $F_z$. By single molecule experiments, Cocco and co-workers showed that the dissociation times associated with the unzipping of DNA helices is a quadratic function of the force in the low-force regime [2]. From Fig. 10, we appreciate a slight decrease of the rupture times versus the applied force at $b_d = 2$ and 3 - which is, however, associated with a large uncertainty.

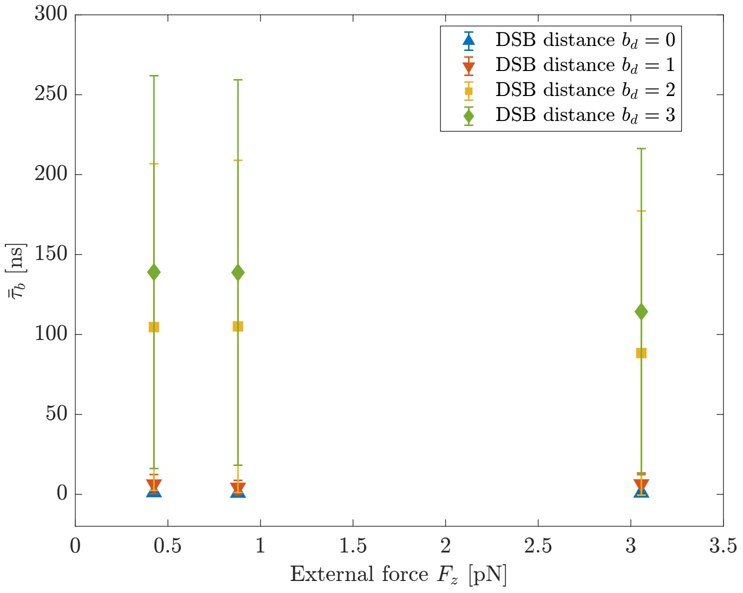


**Supplementary Figure 10:** Zero-force extrapolation of the characteristic rupture times of DNA by the diverse DSB motifs, reporting of a negligible effect of $F_z$ in the low-force regime.
